# An optimised protocol for isolation of RNA through laser capture microdissection of leaf material

**DOI:** 10.1101/644997

**Authors:** Lei Hua, Julian M Hibberd

## Abstract

Laser Capture Microdissection is a powerful tool that allows thin slices of specific cells types to be separated from one another. However, the most commonly used protocol, which involves embedding tissue in paraffin wax, results in severely degraded RNA. Yields from low abundance cell types of leaves are particularly compromised. We reasoned that the relatively high temperature used for sample embedding, and aqueous conditions associated with sample preparation prior to microdissection contribute to RNA degradation. Here we describe an optimized procedure to limit RNA degradation that is based on the use of low melting point wax as well as modifications to sample preparation prior to dissection, and isolation of paradermal, rather than transverse sections. Using this approach high quality RNA suitable for down-stream applications such as quantitative reverse transcriptase polymerase chain reactions or RNA-sequencing is recovered from microdissected bundle sheath strands and mesophyll cells of leaf tissue.

## Introduction

Multicellularity has evolved repeatedly across the tree of life, and is a defining feature of land plants. Not only does multicellularity solve size and lifespan limitations caused by diffusion and ageing of individual cells respectively, it also allows increased complexity through the differentiation of cell types that become specialized for particular functions. Thus, to understand how a multicellular organism is built and then its structures maintained, analysis of its constituent cell types is desirable.

Various methods have been developed to isolate and study specific cell types in plants. In some cases, different tissue types can be separated relatively easily. For instance, in some plant species with C_4_ leaf anatomy bundle sheath strands can be separated from the adjoining mesophyll by differential grinding (Edwards and Black 1971; Kanai and Edwards 1973; Sheen 1995; Covshoff et al. 2013). However, this is not possible in leaves of species that use the far more prevalent C_3_ pathway, or for tissues in other plant organs, and so more complex approaches have been developed. Many of these rely on producing transgenic lines in which a cell autonomous reporter marks a specific cell type such that it can be purified for analysis. This can involve marking cells with a fluorescent protein to allow fluorescence activated cell sorting (Adrian et al. 2015), or placing an exogenous tag onto ribosomes (Mustroph et al. 2009; Aubry et al. 2014) or nuclei (Deal and Henikoff 2011; Sijacic et al. 2018) such that they can be immunopurified and mRNAs sequenced and quantified. The latter approach has been particularly successful in roots where the protoplasting required is relatively fast (Birnbaum et al. 2003; Brady et al. 2007; Li et al. 2016).

However, it is not always possible to generate transgenic plants, or identify a promoter that drives strong expression in the cell type being studied. In the case of leaves, the process of protoplasting is known to generate a significant stress response and de-differentiation (Sawers et al. 2007) such that this approach is compromised if the aim is to better understand photosynthesis. In principle, laser capture microdissection provides an orthogonal method to these approaches, enabling highly purified cell populations to be harvested without requiring the generation of transgenic lines (Nelson et al. 2006). The success of laser capture microdissection largely relies on sample preparation. For example, thin sections need to be produced, but during fixation, embedding and then sectioning, good morphological preservation is required for specific cell types to be dissected. At the same time RNA quality need to be maintained. Freezing and cryosectioning preserve RNA and metabolite composition, but destroy histological details and so have been used in only limited plant species and tissue (Kerk et al. 2003; Nakazono et al. 2003). Chemical fixation followed by paraffin embedding is the most commonly used approach for laser capture microdissection of plant tissue, and so has been used to study cell types from leaves of rice and maize, as well as tomato fruit, soybean roots and Arabidopsis flowers (Klink et al. 2005; Wuest et al. 2014; Jiao et al. 2009; Gandotra et al. 2013; Kerk et al. 2003; Aubry et al. 2014; Aubry et al. 2016). Typically, in these studies non-crosslinking solutions such as Farmer’s fixative or acetone are used to stabilize RNA, and the dehydrated tissue is then mounted in paraffin wax at ∼60°C to allow thin sections to be subjected to microdissection. Although histological details are well preserved using this method, considerable RNA degradation can take place (Gomez et al. 2009; Roux et al. 2018). We found this to be a particular problem with low abundance cell types of leaves. To address this issue, we sought to modify existing protocols to increase RNA yield and integrity during sample processing as well as the laser capture microdissection procedure itself. By adopting a low-melting point wax, as well as modifying sample preparation prior to microdissection and isolation of paradermal rather than transverse sections, we provide a simple and robust method to allow high quality RNA to be obtained from specific cells of leaves that are not accessible using existing methodologies.

## Materials and Methods

### Plant materials and growth

Seeds of *Arabidopsis thaliana* ecotype Columbia were sown in 1:1 mixture of Levington M3 high nutrient compost and Sinclair fine Vermiculite soil, vernalized for 3 days and then transferred to a controlled environment room set at 22°C with a photoperiod of 16h light and 8h dark, a photon flux density of 200 μmol photons m^-2^ s^-1^. Rice (*Oryza sativa* ssp. *indica* IR64) was germinated and grown in 1:1 mixture of top soil and sand for two weeks in a controlled environment growth room set at 28 °C day 25 °C night, a relative humidity of 60%, a photoperiod of 12h light and 12h dark, and a photon flux density of 300 μmol m^-2^ s^-1^.

### Sample preparation

To evaluate the effect of fixative on RNA integrity, fully expanded leaves of Arabidopsis or rice were sampled and fixed in ice-cold 100% (v/v) acetone or Farmer’s fixative (75% (v/v) ethanol, 25% (v/v) acetic acid) for 2 hours and 4 hours on ice, respectively, prior to immediate RNA extraction. To conduct laser capture microdissection rice leaves were cut into 5-8 mm pieces with RNAZap treated scissors, and fixed under vacuum for two 10 minutes periods in ice-cold 100% (v/v) acetone and then left with gentle stirring for 3 hours. Arabidopsis leaves were treated in the same way, but to maintain tissue structure not subjected to vacuum infiltration. Leaf tissue was then dehydrated through an ice-cold series of 70%, 85%, 95% and 100% (v/v) ethanol for 1 hour each. Samples were incubated in 100% (v/v) ethanol overnight at 4°C, prior to being placed in 25%, 50%, 75% and then 100% Steedman’s wax at 37°C for 2 hours. This final solution of 100% Steedman’s wax was replaced twice every 2 hours. Tissue was embedded in a 9-cm petri-dish, and after wax had solidified it was cut into 1 cm^3^ blocks and stored in 50 ml falcon tubes with self-indicating silica gel at -80°C. Steedman’s wax was prepared as described (Vitha et al. 2000), 1000 g polyethylene glycol 400 distearate and 111 g 1-hexadecanol were melted at 60 °C and mixed thoroughly, prior to being aliquoted into 50 ml RNase-free Falcon tubes. Tissue embedded in paraffin wax was also processed on ice, and dehydration and embedding took place in an automated embedding machine that moved samples though a series of 50%, 70%, 95% and 100% (v/v) ethanol, followed by 4 x 1 hour incubations in 100% (v/v) HistoClear, and 2 x 1 hour incubations in Paraplast plus at 60 °C under vacuum.

### RNA extraction

RNA extraction from whole tissues was undertaken using the RNeasy Plant Mini Kit (Qiagen), and from microdissected cells using the PicoPure™ RNA Isolation Kit with on-column DNaseI treatment. To quantify yields 1.5 μl samples of eluted RNA were denatured in 0.2 ml RNase-free tubes at 70°C for 2 minutes. and 1 μl was analysed using an Agilent Bioanalyser RNA 6000 Pico assay. Electropherograms were assessed qualitatively for background signal, and the appearance of cytosolic and chloroplastic ribosomal RNA peaks, and quantitatively using the common metrics of the 25S to 18S ribosomal RNA ratio, and the RNA Integrity Number (RIN) (Schroeder et al. 2006).

### Sectioning and laser capture microdissection

Paradermal sections were prepared using a microtome. Paraffin embedded sections were placed onto a dry polyethylene naphthalate (PEN) membrane slide (Arcturus) and then floated on diethyl pyrocarbonate (DEPC)-treated water at 42°C to expand the sections and ensure they were flat. Water was then removed and slides dried at 42°C for 20 min. Steedman’s wax embedded sections were similarly expanded on DEPC-treated water at room temperature on a PEN membrane slide, before the slide was dried using tissue paper at room temperature. Before laser capture microdissection, paraffin wax was removed by incubating slides in 100% (v/v) Histoclear for 5 minutes, whilst Steedman’s wax was removed by incubating slides in 100% (v/v) acetone for 1 minute at room temperature. Laser capture microdissection was performed on an Arcturus Laser Capture Microdissection system using Capsure macro caps to collect bundle sheath strands and mesophyll cells.

## Results and Discussion

### RNA integrity is maintained with non-crosslinking fixatives

To limit RNA degradation, tissue fixation needs to be rapid. Compared with cross-linking fixatives, precipitative fixatives such as acetone and Farmer’s fixative have commonly been used for laser capture microdissection sample preparation because they retain good histological detail as well as reasonable RNA yields (Kerk et al. 2003). However, to our knowledge, a quantitative analysis of the effect of these fixatives on RNA integrity has not been reported. We therefore fixed Arabidopsis leaves using acetone or Farmer’s fixative on ice for 2 and 4 hours, extracted RNA and found that yield and integrity were similar after 2 hours and 4 hours fixation using either fixative (Supplemental Figure 1). This suggests that RNA was preserved well by each of these precipitative fixatives. However, it was noticeable that leaf tissue sank more rapidly in acetone than in Farmer’s fixative, suggesting a faster penetration into leaf tissue. We therefore subsequently used acetone for sample preparation.

**Figure 1:**
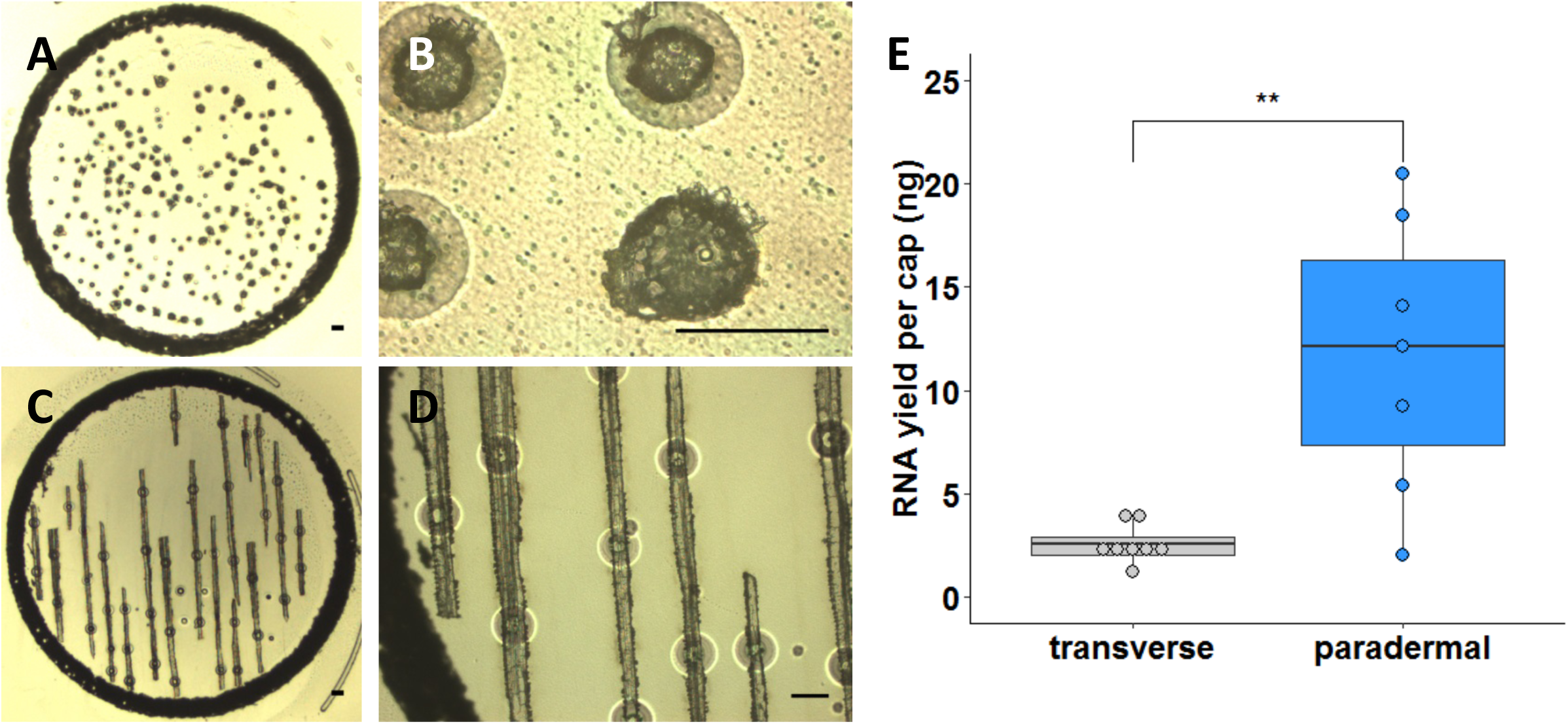
Isolation of RNA from paradermal sections increases yield from Bundle Sheath Strands (BSS). Representative images of BSS dissected from transverse **(A,B)** or paradermal sections **(C,D)** of rice leaves. **A** and **C** show an entire cap used to collect samples after laser capture microdissection. **B** and **D** show higher magnification images of individual BSS in transverse **(B)** or paradermal section **(D)** respectively. Scale bars represent 100 µm. **(E)** Quantitation of RNA yield after microdissection of BSS tissue from transverse or paradermal sections. Data are derived from 7 or 8 biological replicates, and were subjected to a one tailed t-test, where ** P<0.01.

### RNA integrity is improved after Steedman’s wax infiltration

The most commonly used embedding medium for laser capture microdissection studies of plants is paraffin wax, presumably due to its ease of handling and good preservation of histological details. Therefore, initially we embedded rice leaves using paraffin wax and transverse sections were prepared to isolate mesophyll and bundle sheath strands (Figure 1A&B). However, even when a cap was fully loaded with tissue, which takes around 2 hours of continuous microdissection, very low quantities of RNA were obtained from bundle sheath strands (Figure 1E). We therefore tested whether sampling from paradermal sections (Figure 1C&D) improved yields. About ten paradermal sections could be prepared from one leaf, and in approximately one hour, nearly all of the bundle sheath strands in these sections could be dissected. This yielded significantly greater amounts of RNA (Figure 1E). Thus, paradermal sectioning resulted in more tissue being captured per slide (Figure 1A and 1C), was about twice as quick, and so reduces the risk of RNA degradation. However, consistent with reports on other tissues (Roux et al. 2018), we also found that RNA quality from paraffin embedded tissue was low. Since the fixation process itself appeared not to have a deleterious effect on RNA quality (Supplemental Figure 1), we reasoned that losses in RNA integrity were caused by fragmentation occurring at the relatively high temperatures associated with infiltration of paraffin wax. We therefore identified Steedman’s wax as an alternative, low-melting point embedding medium, which has been used in immuno-localization experiments in animals, as well as laser capture microdissection studies of roots, nodules and embryos (Steedman 1957; Vitha et al. 2000; Gomez et al. 2009; Thiel et al. 2011; Limpens et al. 2013, Roux et al. 2014).

To test the applicability of Steedman’s wax for leaf tissue, it was used to embed Arabidopsis and rice leaves, RNA was extracted from whole microtome sections (without mounting on slides) and the yield and integrity compared with that recovered from equivalent paraffin-embedded sections. RNA isolated from tissue in Steedman’s wax showed less elevated baselines and clearer peaks representing the cytosolic and chloroplastic ribosomal RNAs (Figure 2A-D). Both the RNA Integrity Number (RIN) and ribosomal 28S:18S RNA ratio were statistically significantly higher when Steedman’s wax was used to embed Arabidopsis leaves (Figure 2E&F). Although the RIN values from rice leaves were not increased significantly, it was noticeable that the ribosomal RNA peaks were more defined, and that the ratio of the cytosolic ribosomal 25S to 18S RNAs was significantly higher when Steedman’s embedding medium was used (Figure 2E&F). Taken together, these findings indicate that RNA recovered from leaves embedded in Steedman’s wax was of higher quality than that isolated after embedding in paraffin wax. Whilst we are not able rule out other effects, the simplest explanation is that the lower temperature used during infiltration of Steedman’s wax leads to less RNA damage.

**Figure 2:**
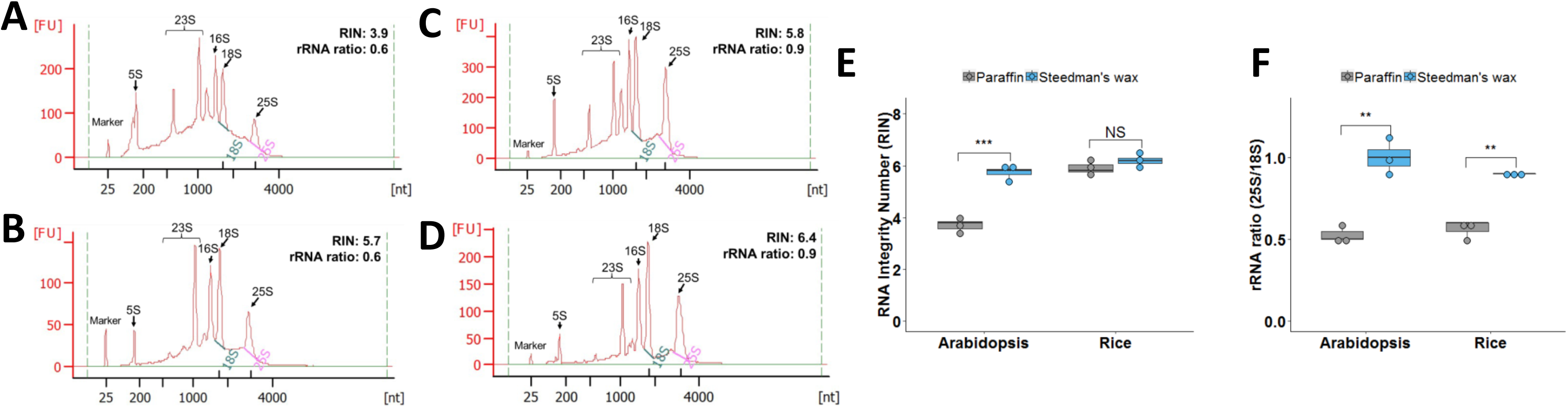
Steedman’s wax improves RNA integrity from sections of Arabidopsis and rice. Representative RNA profiles of Arabidopsis leaves embedded using paraffin **(A)** or Steedman’s wax **(B)**. Representative RNA profiles of rice leaves embedded using paraffin **(C)** or Steedman’s wax **(D)**. The major ribosomal peaks are annotated. The y-axis shows Fluorescence Units (FU) and the x-axis nucleotide length. Quantitation of RNA Integrity Number **(E)** and rRNA ratio **(F)** after embedding in paraffin or Steedman’s wax. Both the RNA Integrity Number (RIN) and rRNA ratios were higher when samples were embedded in Steedman’s wax. Data were subjected to a one tailed t-test, where NS = Not Statistically different, ** P<0.01, *** P<0.001.

To determine whether the duration of infiltration in Steedman’s wax affects RNA quality, we extracted RNA from rice leaf tissue after 1 hour, 3 hours, or 6 hours of infiltration in Steedman’s wax. Both the RIN value and ribosomal 28S to 18S RNA ratio remained essentially unchanged over this time-course, suggesting that in species that require longer infiltrations for good sectioning, extending the infiltration time in wax could be used without compromising RNA quality (Supplemental Figure 2).

### A procedure to minimize RNA degradation during slide preparation

Subsequent to wax embedding, but prior to laser capture microdissection, there are further opportunities for RNA to be degraded. For example, during slide preparation, sections are typically expanded by floating on warm RNase-free water at 42°C to ensure they lie flat on the microscope slides. Water is then removed and samples dried at 42°C for 20-30 minutes. Consistent with RNA degradation during this process, after slide preparation from paraffin-embedded rice tissue the cytosolic and chloroplastic ribosomal RNA peaks were less defined, yields were lower, and the 28S to 18S ribosomal RNA ratio was lower compared with to that isolated from freshly-cut sections (Figure 3A-D). In contrast, after slide preparation using Steedman’s wax for embedding, the cytosolic and chloroplastic ribosomal RNA peaks were clearly defined, and the 28S to 18S ribosomal RNA ratio was maintained (Figure 3E-3H). We also found that sections embedded in Steedman’s expanded immediately on water at room temperature (20°C), and that the water could be removed rapidly by absorption onto soft tissue paper. Adhesion of thin sections to the slide was enhanced by providing gentle pressure with dry tissue paper (Supplemental Figure 3). This rapid process avoids the prolonged exposure of sections to higher temperatures and so preserves RNA integrity during slide preparation. Examination of tissue integrity using light microscopy after embedding in Steedman’s wax showed that histological details were as good as those seen after embedding in paraffin wax (Supplemental Figure 4).

**Figure 3:**
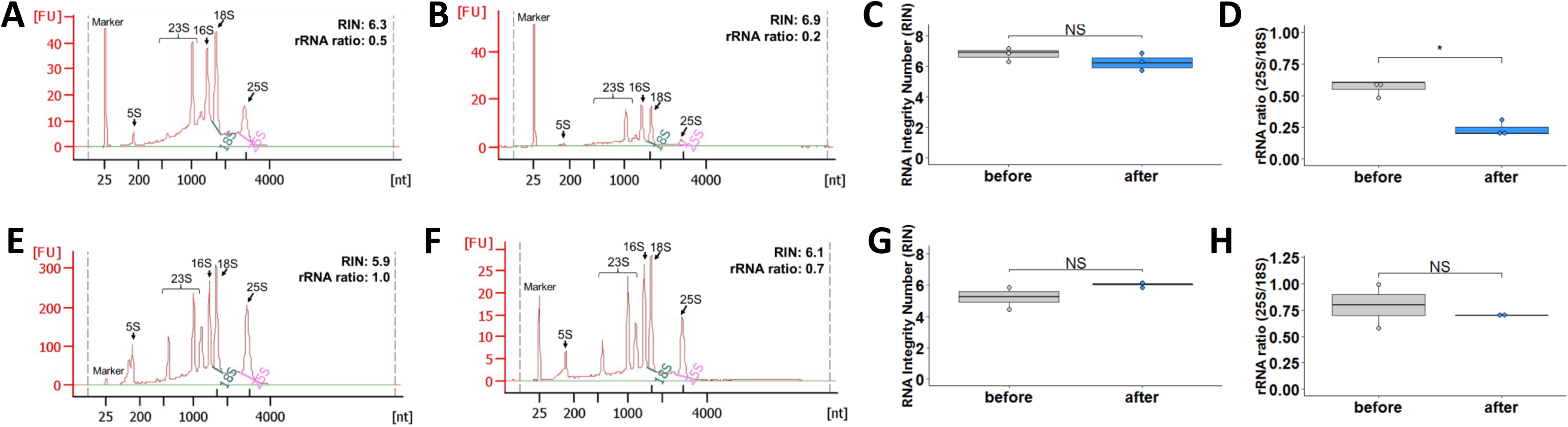
Limited degradation of RNA from Steedman’s wax embedded sections occurs when slide preparation is optimised. **(A&B)** Representative RNA profiles of rice sections embedded in paraffin before **(A)** and after **(B)** slide preparation. Comparison of RIN **(C)** and rRNA ratio **(D)** of paraffin embedded sections before and after slide preparation. **(E&F)** Representative RNA profiles of rice sections embedded in Steedman’s wax before **(E)** and after **(F)** slide preparation. Comparison of RIN **(G)** and rRNA ratio **(H)** of Steedman’s wax embedded sections before and after slide preparation. In panels A,B,E&F, the major ribosomal peaks are annotated, the y-axis shows Fluorescence Units (FU) and the x-axis nucleotide length. Data were subjected to a one tailed t-test, where NS = Not Statistically different, * P<0.05.

**Figure 4:**
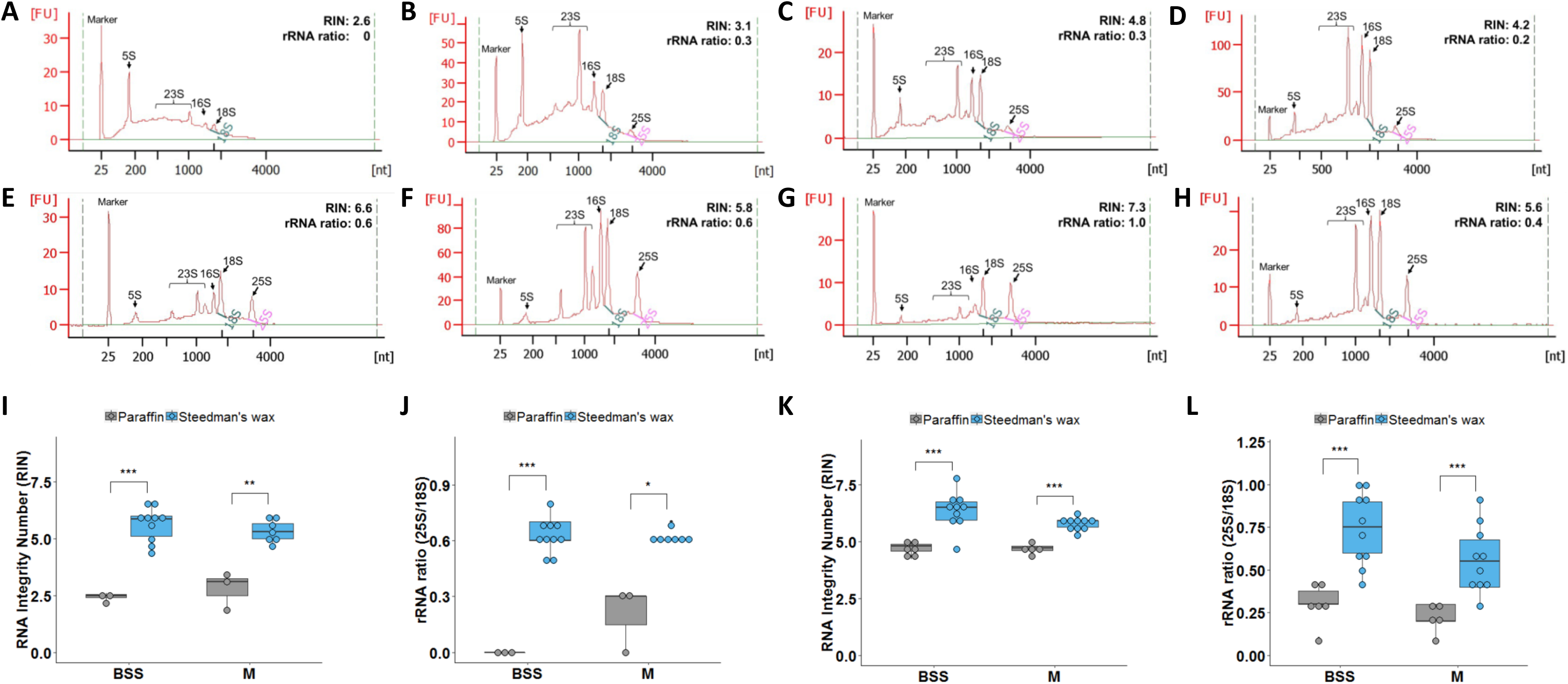
RNA integrity from Bundle Sheath Strands (BSS) and mesophyll cells is significantly improved when Steedman’s wax is used. **(A-D)** Representative RNA profiles of microdissected BSS **(A, C)** or M cells **(B, D)** from Arabidopsis **(A, B)** and rice **(C, D)** embedded in paraffin. **(E-H)** Representative RNA profiles of microdissected BSS **(E, G)** or M cells **(F, H)** from Arabidopsis **(E, F)** and rice **(G, H)** embedded in Steedman’s wax. The y-axis shows Fluorescence Units (FU) and the x-axis nucleotide length. Note the lower background in Steedman’s wax. Comparison of RIN values **(I, K)** and rRNA ratio **(J, L)** of RNA extracted from microdissected BSS and M cells from Arabidopsis **(I, J)** and rice **(K, L)** embedded using paraffin and Steedman’s wax. Data were subjected to a one tailed t-test, where * P<0.05, ** P<0.01, *** P<0.001.

### High quality RNA can be obtained after laser capture microdissection

To assess the combined impact of the modifications documented above on the final RNA quality obtained from laser capture microdissection, we compared RNA quality from microdissected tissues embedded in either paraffin or Steedman’s wax. For this purpose, bundle sheath strands and mesophyll cell sections were captured from both Arabidopsis and rice leaf tissues. RNA isolated by laser capture microdissection from paraffin embedded sections of either species showed either no clear, or compromised ribosomal RNA peaks (Figure 4A-D). This was particularly noticeable for the bundle sheath strands. Moreover, the baseline was high (Figure 4A-D) indicating that the RNA was severely degraded, and quantitation confirmed these qualitative assessments (Figure 4I-L). In contrast, RNA isolated from either Arabidopsis or rice tissue embedded in Steedman’s wax showed less elevated baselines, more defined ribosomal RNA peaks (Figure 4E-H), and higher RIN values and 28S to 18S ribosomal RNA ratios (Figure 4I-L).

With the advances and reduced cost of next-generation sequencing, RNA-SEQ has become a common tool for profiling transcript abundance. However, a high quality RNA input is important for reliable and reproducible results. For example, it has been reported that RNA degradation can have a broad effect on quality of RNA-SEQ data, including 3’ bias in read coverage, quantitation of transcript abundance, increased variation between replicates, and reductions in library complexity (Chen et al. 2014, Feng et al. 2015, Gallego et al. 2014). Indeed, Gallego et al. (2014) found that RIN values are a robust indicator for RNA degradation, and that RNA-SEQ data from RNA with RIN values >5 show better correlation with intact RNA. For cell specific profiling of gene expression using laser capture microdissection, the RIN value of microdissected RNA was rarely >5 when paraffin wax was used during sample preparation. In contrast, our optimised sample preparation method using low-melting temperature wax, led to most RNA isolated after microdissection having RIN values >5. We therefore conclude that these simple modifications allow tissue to be prepared such that different cell types in the leaf can still be identified, and that the quality of the RNA available for sampling is improved. We anticipate that this approach will greatly facilitate the analysis of gene expression in specific cell types of leaves.

## Acknowledgments

The work was supported by a C_4_ Rice project grant from The Bill and Melinda Gates Foundation to the University of Oxford (2015-2019).

## Author contribution

Conceptualization, Lei Hua and Julian M Hibberd; Data curation, Lei Hua and Julian M Hibberd; Formal analysis, Lei Hua; Funding acquisition, Julian M Hibberd; Investigation, Lei Hua; Methodology, Lei Hua; Project administration, Julian M Hibberd; Supervision, Julian M Hibberd; Writing – original draft, Lei Hua; Writing – review & editing, Lei Hua and Julian M Hibberd.

