## Supplemental Figures for "An optimised protocol for isolation of RNA through laser capture microdissection of leaf material"

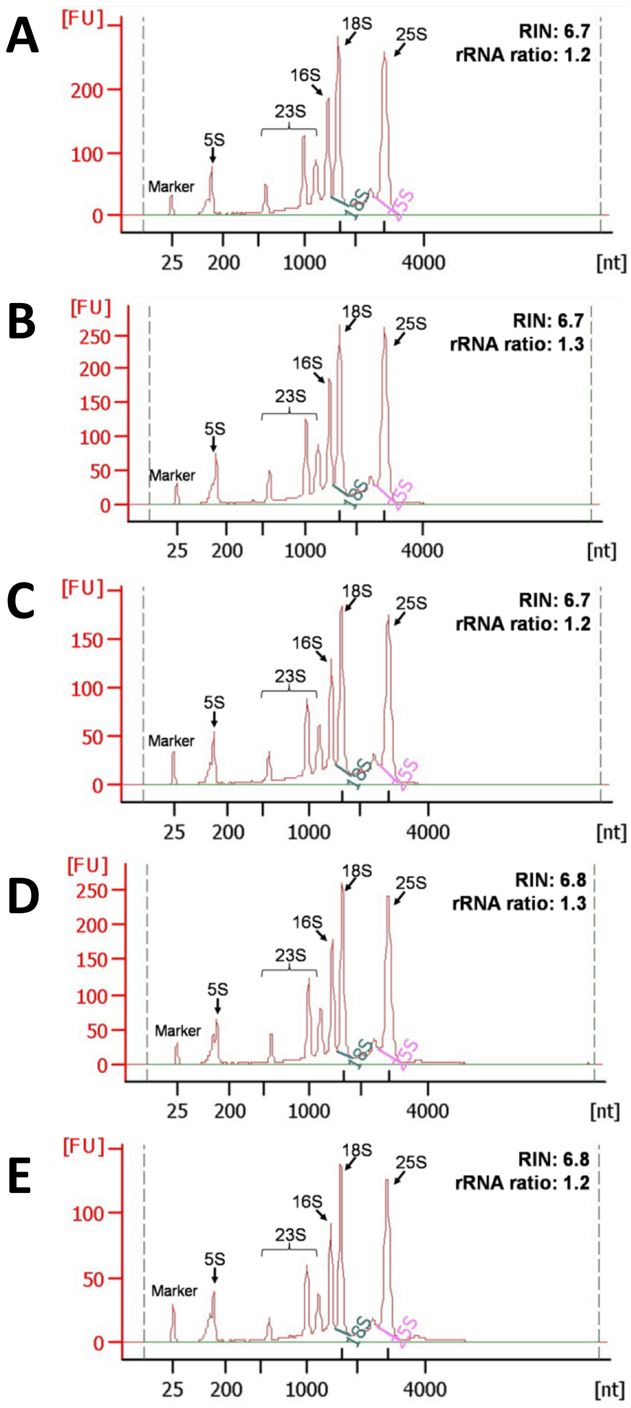


**Supplemental Figure 1: RNA integrity after fixation of leaves with 100% (v/v) acetone or Farmer's fixative.** Bioanalyzer traces derived from RNA extracted from snap-frozen *Arabidopsis* leaves **(A),** leaves placed in 100% (v/v) acetone for after 2 **(B)** or 4 hours **(C)**, and leaves placed in Farmer's fixative for 2 **(D)** or 4 hours **(E)**. The major ribosomal RNA peaks are annotated. The y-axis shows Fluorescence Units (FU) and the x-axis nucleotide length.

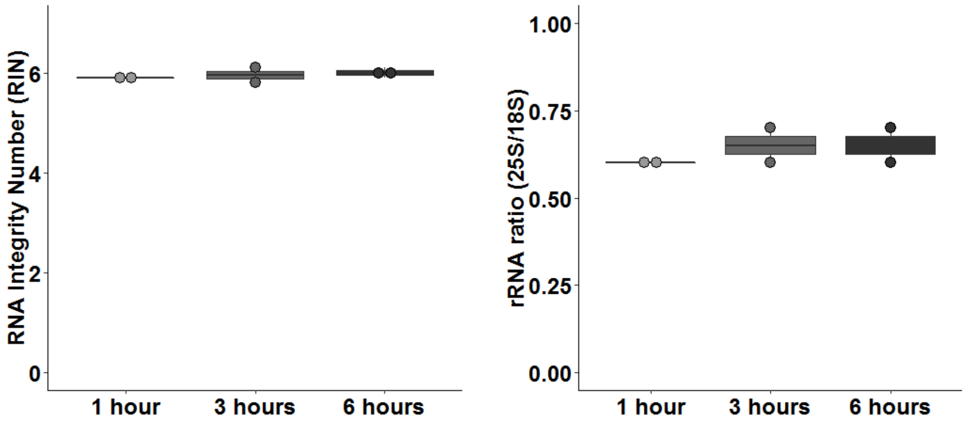


**Supplemental Figure 2: Infiltration with Steedman's wax stabilizes RNA quality.** RIN values (A) and rRNA ratio (B) of RNA from rice leaves after 1 hour, 3 hours, and 6 hours of infiltration in Steedman's wax at 40 ˚C.

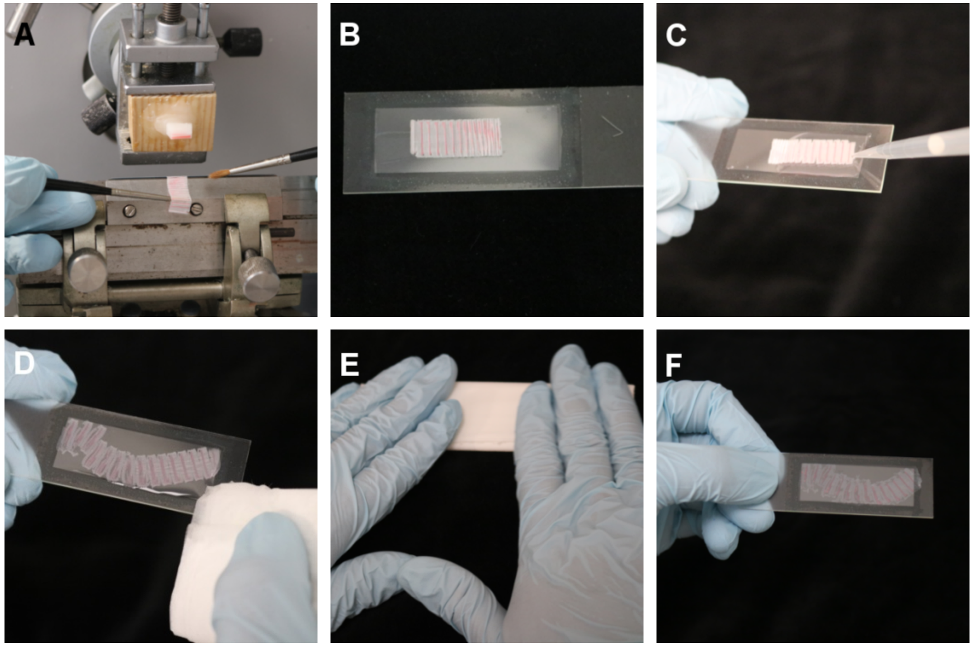


**Supplemental Figure 3: Key stages in sample adhesion to slides for tissue embedded in Steedman's wax.** Photographs illustrate the main steps associated with sectioning and slide preparation prior to Laser Capture Microdissection (LCM). **A.** Sectioning of Steedman's wax embedded leaf with microtome. **B.** The wax ribbon is placed on membrane slide. **C.** The wax ribbon is expanded and flattened onto the slide using DEPC-treated water at room temperature. **D.** After ribbon expansion water is removed using tissue paper. **E.** Dry sections are generated by providing gentle pressure on each section with folded tissue paper. **F.** Exemplar slide with leaf tissue that could be used for LCM directly.

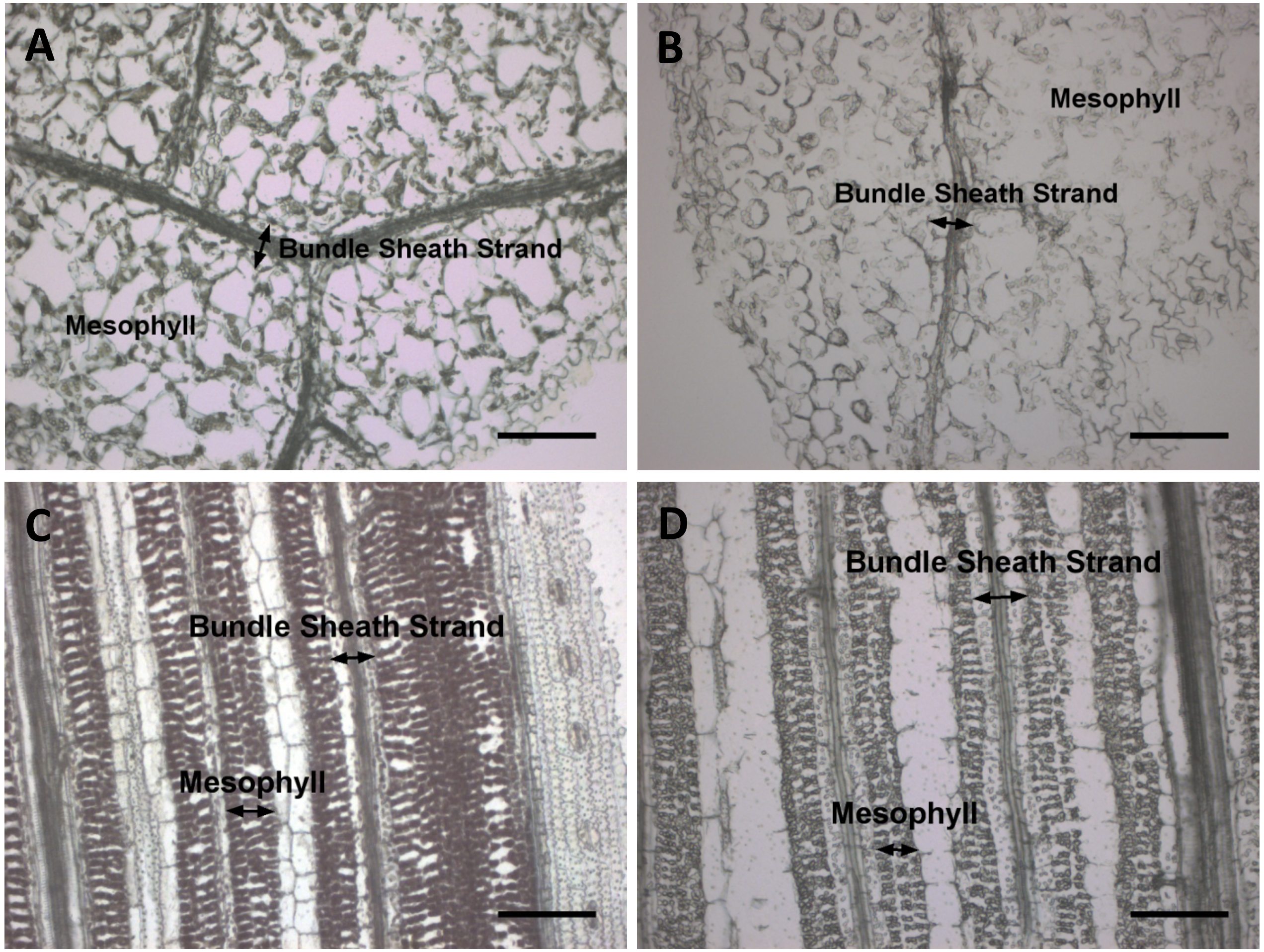


**Supplemental Figure 4: Identification of BSS and M cells in Arabidopsis and rice leaf sections embedded with paraffin (A, C) or Steedman's wax (B, D).** Bundle sheath strands and mesophyll cells are marked with arrows. Scale bars represent 100 µm.
